# Annelid eye evolution revealed by developmental, ultrastructural, and connectome analyses of cerebral eyes in *Malacoceros fuliginosus*

**DOI:** 10.64898/2026.02.04.703583

**Authors:** Suman Kumar, Anna Seybold, Oleg Tolstenkov, Sharat Chandra Tumu, Harald Hausen

## Abstract

Eye evolution has long attracted interest, yet how multiple cerebral eyes within a lineage originate and diversify remains unclear. Annelids display exceptional diversity in eye number and structure, but the homology and function of distinct eye pairs are poorly understood. Here we investigate the cerebral eyes of the sedentary annelid *Malacoceros fuliginosus* using an integrated developmental, molecular, ultrastructural, and connectomic approach. We show that both the early-developing ventral and later-developing dorsal eyes are simple, few-celled, inverted rhabdomeric eyes that express transcription factors with conserved roles in animal eye development. Two r-opsin paralogs and distinct neurotransmitters are differentially expressed in different photoreceptor cells of the eyes. Ultrastructural reconstructions across larval development reveal differences in cellular composition and growth dynamics, while axonal tracing shows that photoreceptors from ventral and dorsal eyes project to overlapping regions of the larval brain. The overall organization and projections resemble those described in the errant annelid *Platynereis dumerilii*. Together, these data support the hypothesis that an ancestral cerebral eye duplicated early in annelid evolution, giving rise to multiple eye pairs with stage-specific functions.

## Introduction

For light-exposed animals, vision plays a key role in navigating and interacting with their surroundings. As a minimum requirement, directional light detection relies on eyes composed of photoreceptive cells and shielding structures that block light from one side. Although opsin based light detection likely evolved only once in animals, the homology and independent origins of various eye types and their components across animal lineages remain debated (Salvini-Plawen & Mayr 1977; Eakin 1979; Arendt 2003; Oakley & Speiser 2015) While molecular profiles point towards conservation and diversification of key components such as photoreceptor and shielding pigment cells, convergent evolution of eye complexity and independent emergence of accessory structures are also supported (Arendt 2003; Ogura et al., 2004; Kozmik et al., 2008; Gehring 2014). The picture is further complicated by the fact that many animals possess multiple pairs of eyes, which may differ in structure and function, raising questions about their specific roles and evolutionary origins.

Among lophotrochozoans, annelids exhibit remarkable diversity in light sensory structures in terms of number, size and organization with both cerebral and ectopic eyes being common among annelids (Eakin & Hermans 1988; Verger-Bocquet 1992; Purschke et al., 2006, 2022). Based on their lifestyle and supported by phylogenomic analyses, annelids are divided into two major groups - Sedentaria and Errantia - and six basal branching groups (Weigert 2016). Across the major annelid groups, both simple and complex eyes occur in varying numbers, whereas in the basal branching taxa, eyes are typically simple and are known from Oweniidae, Chaetopteridae, Amphinomidae and Sipuncula (Purschke et al., 2022). Unlike errant annelids, members of Sedentaria often exhibit structurally simpler eyespots, composed of just pigment cells and photoreceptors, and sometimes lack multiple pairs.

The annelid *Platynereis dumerilii*, a member of Errantia, has emerged as a key model in developmental and sensory biology providing major insights into the function, neural circuitry and evolution of animal eyes (Jékely et al., 2008; Randel et al., 2013, 2014, 2015). Like many species of this group, they are characterized by an active lifestyle, possess complex adult eyes with a multicellular retina and a lens-structure (Rhode 1992). In *P. dumerilii*, larvae bear a pair of simple eyespots in addition to the more complex adult eyes. Both eyes share expression of two paralogs of the r-opsin visual pigment and a conserved set of eye developmental genes, suggesting a close evolutionary relationship (Arendt et al., 2002). However, they differ in their primary neuronal targets and neurotransmitter profile: the simple eyespots are acetylcholinergic and project to ciliated locomotor cells, mediating early phototactic behavior (Jékely et al., 2008), while the glutamatergic adult eyes connect to the brain and control musculature via complex circuits (Randel et al. 2014, 2015). Whether this dual-eye system is an ancestral or derived annelid feature remains unresolved.

To address this, we studied the eye structure, development, molecular features, and eye connectome in *Malacoceros fuliginosus*, a hemi-sessile annelid from the sister clade Sedentaria. Our aim was to determine whether features found in *P. dumerilii* are shared with *M. fuliginosus*, thereby providing insight into ancestral versus derived eye conditions. Of the three eyespots that *M. fuliginosus* possesses two exhibit a microvillar organization typical of annelid cerebral eyes, while the third displays a ciliary morphology - an uncommon trait in annelids. By comparing molecular features including opsin expression and the circuitry of the microvillar eyes on the level of individual cells throughout development with data from *P. dumerilii*, we find striking similarities. We suggest that the ventral and dorsal eyes in both species are homologous, despite the vast complexity of the *P. dumerilii* adult eye. Our findings support the idea that the annelid ancestor of Errantia and Sedentaria possessed two pairs of cerebral eyes, which originated not before the duplication of the visual r-opsin.

## Materials and methods

### *Malacoceros fuliginosus* culture

Adult *Malacoceros fuliginosus* were collected from Pointe de Mousterlin, Fouesnant, France, and were maintained in sediment containing seawater tanks at 18°C. Individual males and females were picked, rinsed several times with filtered seawater, and kept in separate bowls until they spawned. The staging was started from the time gametes were combined in a fresh bowl. Bowls were kept at 18°C under 12:12 hr light-dark cycle, and water was replaced every day or every second day. Larvae were fed with microalga *Chaetoceros calcitrans* from 24 hpf onwards after each water change.

### RNA Seq and transcriptome assembly

Total RNA was extracted from cryofixed samples of various larval stages using the Agencourt RNAdvance Tissue Kit (Beckman Coulter). Library preparation and sequencing were performed by EMBL Genomics Core Facility (Heidelberg, Germany) using cation-based chemical fragmentation of RNA, Illumina Truseq RNA-Sample Preparation Kit, and one lane of 100 bp paired-end read sequencing on Illumina HiSeq 2000. Raw reads were trimmed, and error corrected with Cutadapt 1.2.1, the ErrorCorrectReads tool implemented in Allpaths-LG, and assembled with Trinity. A second assembly including several steps for correction of sequencing errors, chimerism, and elimination of redundancy was generated using the DRAP pipeline (Cabau et al. 2017), followed by gene clustering using Corset (Davidson & Oshlack 2014).

### Opsin tree inference

Opsin sequences from *Malacoceros fuliginosus* transcriptome and representative metazoan opsins (Supplementary Table S1) were retrieved by BLAST searches and further screened for the presence of the PFAM 7tm_1 domain and the conserved Lys296 residue predictive of chromophore binding. Representative opsins from cnidarians were included as outgroups. Sequences were aligned using MAFFT v7.490 with the E-INS-i strategy (Katoh and Standley 2013). The conserved seven-transmembrane (7TM) region was extracted based on the DRY/ERY and NPxxY motifs to focus on functionally conserved domains. Columns with <90% occupancy were removed, and the filtered alignment was re-aligned with MAFFT E-INS-i. Maximum likelihood phylogenetic inference was performed with IQ-TREE v1.6.12 (Nguyen et al., 2015) using the LG+F+R8 model selected by ModelFinder (Kalyaanamoorthy et al., 2017). Branch support was assessed using SH-aLRT and ultrafast bootstrap (UFBoot), with 1,000 replicates each; nodes with SH-aLRT ≥75% and UFBoot ≥85% were considered well supported.

Bayesian inference was conducted with MrBayes v3.2.7a (Ronquist et al., 2012) under the LG+F+Γ6 model (two runs of 5 million generations, 25% burn-in). Convergence was verified by an average standard deviation of split frequencies <0.01. Posterior probabilities ≥0.95 were considered strong support. Trees were rooted using cnidarian opsins (*Nematostella vectensis*) and visualized with ggtree (Yu et al., 2017; R Core Team 2023).

### Gene cloning and probe generation

Gene transcripts were retrieved from the assembled transcriptome by bidirectional blast. Specific primers were designed, and fragments were amplified from either mixed stage or stage-specific cDNA. The PCR fragments were inserted into the pGEM-T-easy vector (Promega) and cloned in Top10 E. coli cells (Invitrogen). DIG/FITC labeled RNA probes were generated by DIG RNA labeling mix (Roche) or using transcription reagents along with DIG-UTP/FITC-UTP (Roche) by SP6 or T7 polymerase (Roche). Gene orthology was inferred by reciprocal blast against GenBank.

### *In-situ* hybridization

Larvae were fixed in 4 % PFA (in 1X PBS, 0.1% Tween20) for 2.5 hours at RT. Larvae 48 h and older were first relaxed with 1:1 MgCl2-seawater for 3-5 min before fixing them in 4 % PFA. Samples were stored at −20 °C in methanol until use. The larvae were rehydrated in series of 75, 50, and 25% methanol in PTW (1X PBS pH 7.4, 0.1% Tween20). Tissue was permeabilized by Proteinase K (100 ng/ml) (30 s for 24 h larvae to 3 min for 5 d larvae), followed by 2x5 min Glycine washes (2 mg/ml). The larvae were then acetylated with 1% triethanolamine (TEA) in PTW for 5 min and washed with 0.5 µl/ml acetic acid in 1% TEA for 5 min. Following this, samples were washed 2x5 min in PTW before post-fixing for 15 min in 4 % PFA (in 1X PBS-Tween20). The post-fixed samples were washed four times in PTW for 5 min each. The larvae were then equilibrated in hybridization solution (Hyb solution: 50% formamide, 5X SSC, 50 µg/ml heparin, 100 µg/ml salmon sperm DNA, 0.1% Tween20, 1% SDS and 5% dextran sulfate) for 10 min. Next, samples were prehybridized with Hyb solution at 65 °C for 2-4 h before hybridization with labeled RNA probes at a concentration of 1 ng/µl to 2.5 ng/µl for 48-60 h at 65 °C. The samples were then subjected to two post-hybridization washes of 5 min and 20 min with Hyb solution at 65 °C. The next washes with SSC were done as following: 2X SSC-Hyb solution in series of 25, 50, 75, and 100% 2X SSC each for 10 min at 65 °C, followed by two 30 min 0.05X SSC washes at 65 °C. The samples were equilibrated at RT for 10 min before washing in PTW-0.05X SSC in a series of 25, 50, 75, and 100% PTW for 5 min each. Next, the samples were blocked in Roche Blocking solution (in Maleic acid buffer pH 7.5) for 1 h and then incubated in anti-DIG-AP (Roche) or anti-FITC-AP (Roche) Fab fragments (1:5000) overnight at 4 °C. After this, the larvae were washed six times in PTW for 1 h and then equilibrated first in Mg2-free AP buffer (two 5 min washes), then in AP buffer (100 mM NaCl, 50 mM MgCl2, 100 mM Tris (pH 9.5 for NBT/BCIP staining and pH 8.2 for Fast Blue/Fast Red staining), 0.1 % Tween20) (2x5 min washes). Probe detection was performed using NBT/BCIP (Roche) in AP buffer (pH 9.5) by adding 2.25 µl/ml NBT (from 100 mg/ml stock) and 3.5 µl/ml BCIP (from 50 mg/ml stock). Fast Blue (Sigma)/Fast Red (Roche) double staining was according to the protocol described in Lauter et al., 2011.

### Immunolabeling

Larvae were fixed in 4 % PFA (in 1X PBS, 0.1% Tween20) for 30 min at RT. Larvae 48 h and older were first relaxed with 1:1 MgCl2-seawater for 3-5 min before fixing them in 4 % PFA. After fixing, the samples were washed (2x5 min) in PTW followed by (4x5 min) THT washes (0.1 M Tris pH 8.5, 0.1% Tween20). Samples were then blocked in 5% sheep serum in THT for 1 h before incubating in primary antibodies (r-opsin1 and r-opsin3, 1:100; monoclonal anti-acetylated α-tubulin, 1:300 Sigma, T7451) for 48h at 4°C. The samples were then subjected to 2x10 min washes in 1 M NaCl in THT followed by 5x30 min washes in THT before incubating in secondary antibodies (Alexa Fluor 1:500, Thermo Fisher Scientific) overnight at 4°C. Next, the samples were washed in 2x5 min THT, followed by 5x30 min washes. Specimens were stored in embedding medium (90% glycerol, 1x PBS, and 1% DABCO) at 4 °C until imaging.

### Microscopy and image processing

Light microscopy images were acquired using the Zeiss Examiner A1 microscope. Fluorescence images were obtained using a Leica SP5 confocal microscope. Image processing was performed using ImageJ, Adobe Photoshop CS6, Imaris (Bitplane), and final figures were assembled using Adobe Illustrator CS6.

### Electron microscopy

For electron microscopy, larvae were processed as described in (Seybold et al., 2025) with minor modifications for 72-hpf larvae. In brief, larvae were relaxed for 5 min in 7% MgCl2 and seawater mixed 1:1 and then fixed in 2.5% glutaraldehyde in PBS, post-fixed in 1% osmium tetroxide in the same buffer, (72-hpf larvae were en-bloc stained with reduced Osmium), dehydrated in a graded acetone series and embedded in Epon/Araldite. All steps were performed in a microwave oven (PelcoBioWave^®^Pro +, Ted Pella, Redding, CA, USA). Serial sections of 50 nm (70nm for 72-hpf larvae) were cut with an ultra 35° diamond knife (Diatome, Biel, Switzerland) and a UC7 ultramicrotome (Leica), mounted on PEI coated Beryllium-Copper slot grids (Synaptek, Reston, VA, USA) and contrasted with 2% uranyl acetate and lead citrate. Sectioning was performed longitudinally from dorsal to ventral. For early stages, a complete series with 613 slices (22 hpf) and 619 slices (28 hpf) with an average FoV of 200×100 μm was imaged through the entire thickness of the larvae, with a total section loss of approximately 3% in both stages. For later stages, the anterior half of the larvae was imaged with a slightly higher section loss. Sections were imaged at 4 nm/pixel resolution with a STEM detector in a Supra 55VP (Zeiss, Oberkochen, Germany) equipped with Atlas (Zeiss) for automated large field of view imaging. Image stacks were further processed with Adobe Photoshop CC, chopped to 4096 × 4096 via Matlab, and then registered rigidly followed by affine and elastic alignment (Saalfeld et al., 2012) with TrakEM2 (Cardona et al., 2012) implemented in Fiji (RRID:SCR_002285).

## Results

### (a) Duplication of r-opsin within annelids

R-opsins are an ancient class of opsins that was already present in the last common ancestor of bilaterians (Porter et al., 2012; Hathaway et al., 2013). Early in their evolutionary history, r-opsins diverged into two major subgroups: canonical r-opsins and noncanonical r-opsins (Ramirez et al., 2016) In annelids and many other protostomes, canonical r-opsins function as the main visual opsins (Fain et al., 2010; Porter et al., 2012; Hathaway et al., 2013). While canonical r-opsins diverged several times into many paralogs within arthropods, only a single copy is known from most other protostome invertebrates. In the errant polychaete, *P. dumerilii,* two canonical r-opsins (r-opsin1 and r-opsin3) were discovered, both of which are expressed in the cerebral eyes (Randel et al., 2013) To determine, when canonical r-opsins diverged in annelid evolution, we performed a phylogenetic analysis of annelid canonical r-opsins (Fig. 1). The phylogenetic analysis was based on publicly available and our sequences including *Owenia fusiformis*, which represents a basally branching annelid lineage (Weigert et al. 2014), and from *M. fuliginosus*, a member of Sedentaria, the sister group of Errantia that includes *P. dumerilii*. Canonical annelid r-opsins formed two well-supported clades (Fig. 1; Sup. Fig. 2). Clade A comprised r-opsin1 and r-opsin3 from *P. dumerilii* and corresponding paralogs from *M. fuliginosus*. A single r-opsin from *O. fusiformis* grouped with the r-opsin1 subclade in the maximum likelihood tree; however, analysis of functionally critical residues (Sup. Fig. 3) showed that it lacks several diagnostic substitutions distinguishing r-opsin1 from r-opsin3 in crown-group annelids and instead displays intermediate or distinct character states at key positions. This pattern is consistent with retention of an ancestral, pre-duplication r-opsin. The presence of distinct r-opsin1 and r-opsin3 orthologs in both Sedentaria (*M. fuliginosus*) and Errantia (*P. dumerilii*) indicates that the duplication occurred in their last common ancestor, after divergence of basal annelids. In contrast, leeches and *Capitella teleta* retain only a single r-opsin1/3 representative, suggesting secondary loss following duplication (Fig. 1). A second annelid-specific r-opsin clade (Clade B) was also recovered, comprising additional paralogs from *P. dumerilii*, *M. fuliginosus*, and *O. fusiformis*. Conserved r-opsin sequence motifs were retained across both clades (Sup. Fig. 3). These paralogs were not analyzed further, and subsequent analyses focused on the canonical r-opsin1 and r-opsin3 lineage.

**Figure 1:**
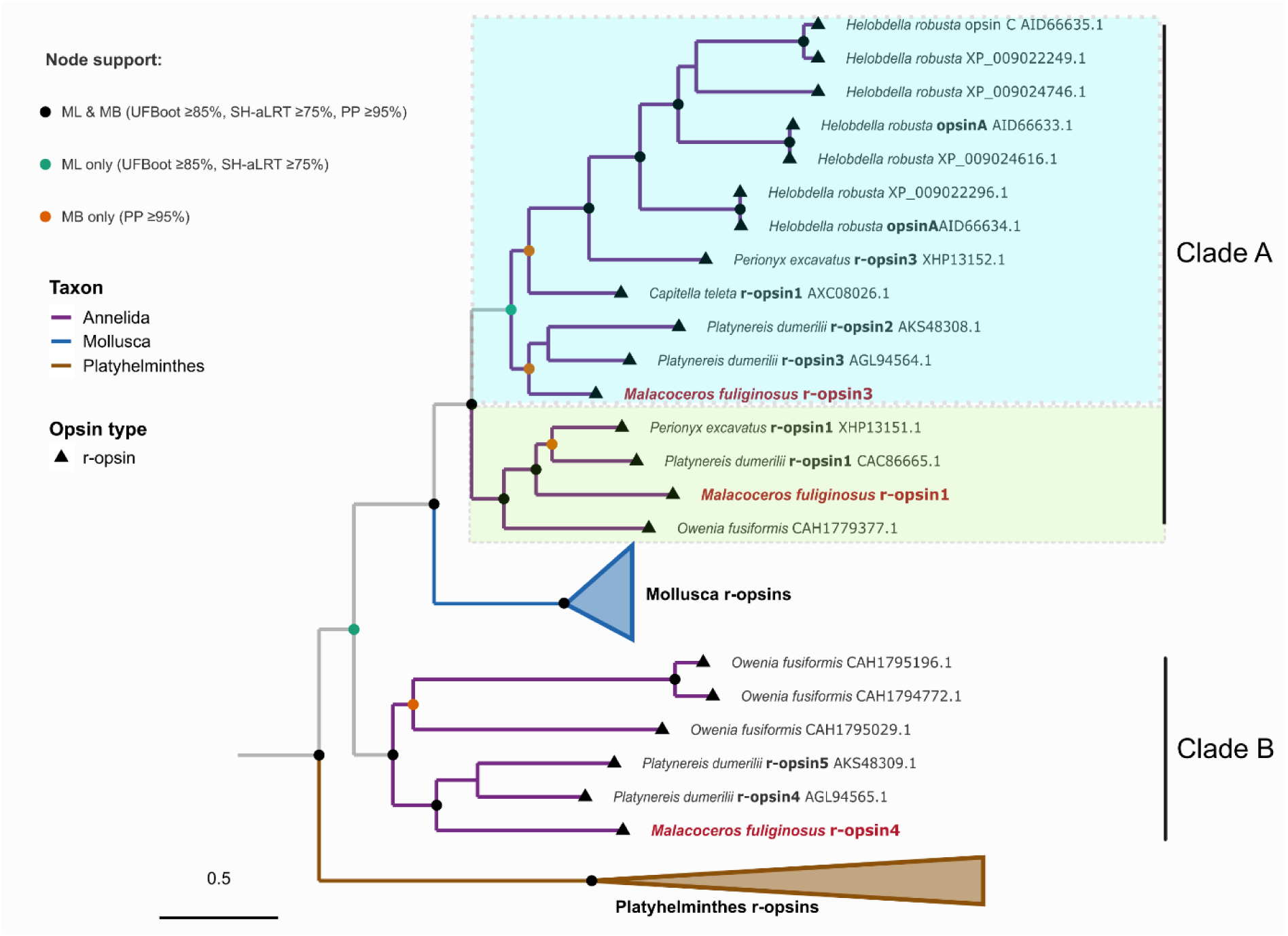
Phylogenetic tree of annelid r-opsins. Maximum-likelihood and Bayesian phylogenetic analysis of r-opsin protein sequences from annelids, along with mollusc and platyhelminth r-opsins. Branch colours indicate taxonomic affiliation: Annelida (purple), Mollusca (blue), and Platyhelminthes (brown). Node support is indicated by coloured circles. Molluscan and platyhelminth r-opsins are collapsed for clarity. Annelid r-opsins resolve into two major, well-supported clades (Clade A and Clade B), consistent with an ancient duplication predating annelid diversification. Canonical r-opsin split within the annelids as indicated by the two groups – r-opsin1 and r-opsin3. *M. fuliginosus* r-opsin1 and r-opsin3 along with the less characterized r-opsin4 is highlighted.

**Figure 2.**
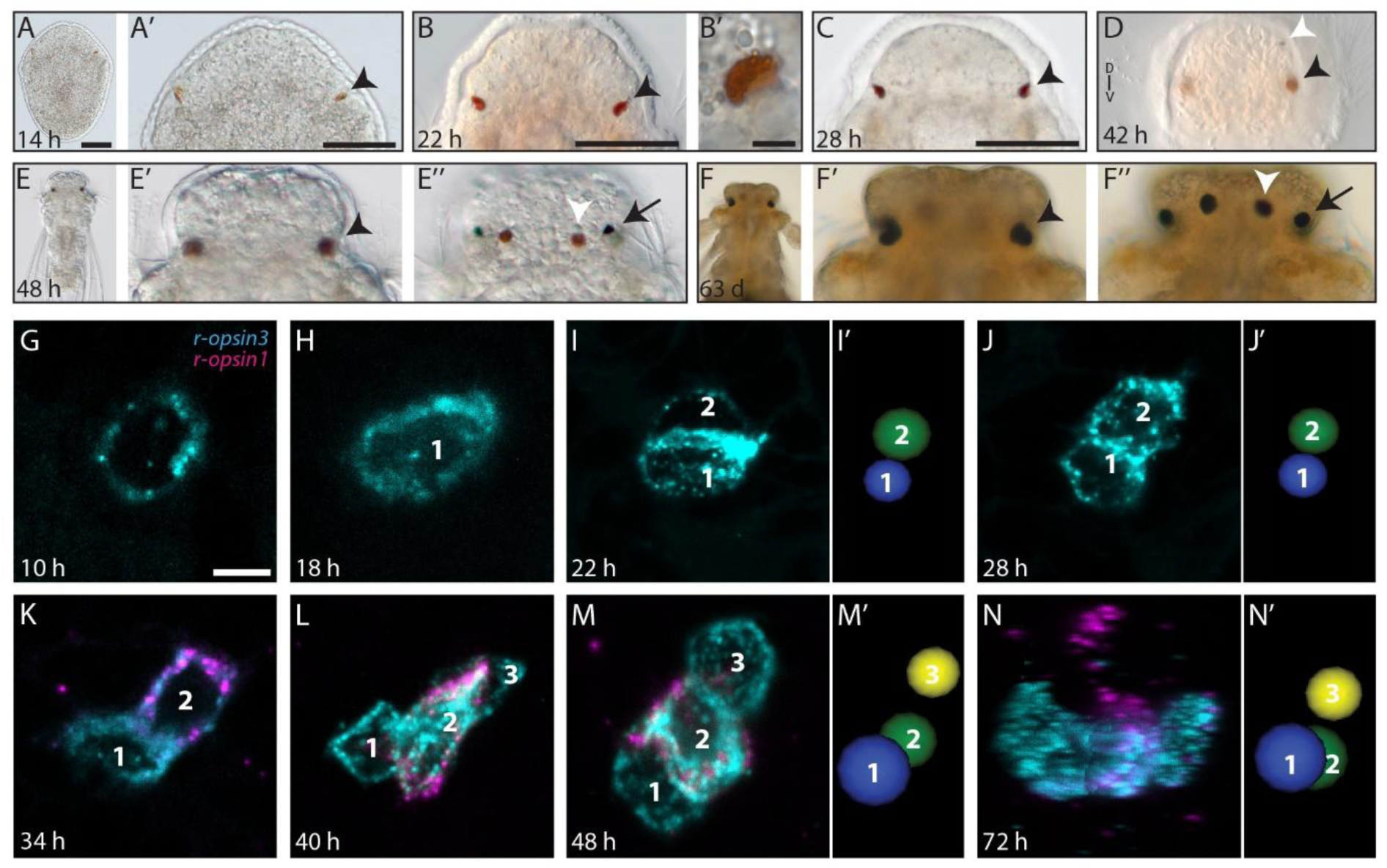
*M. fuliginosus* larval eyes at different developmental stages. (A-F’’) Dorsal overviews and details of the ventral eyes (black arrowhead). (D) Anterior view showing ventral eyes and the right dorsal eye (white arrowhead). (E-F’’) Dorsal views of the ventral eyes, the dorsal eyes and the lateral eyes (black arrow). (G-N’) Expression of r-opsin3 (cyan) and r-opsin1 (magenta) in the PRCs of the right ventral eye at different stages. (I’, J’, M’, N’) 3D view of nuclei locations of PRC1 (blue), PRC2 (green) and PRC3 (yellow). Scale bars: A, A’, B, C: 50 µm; others: 5 µm.

**Figure 3.**
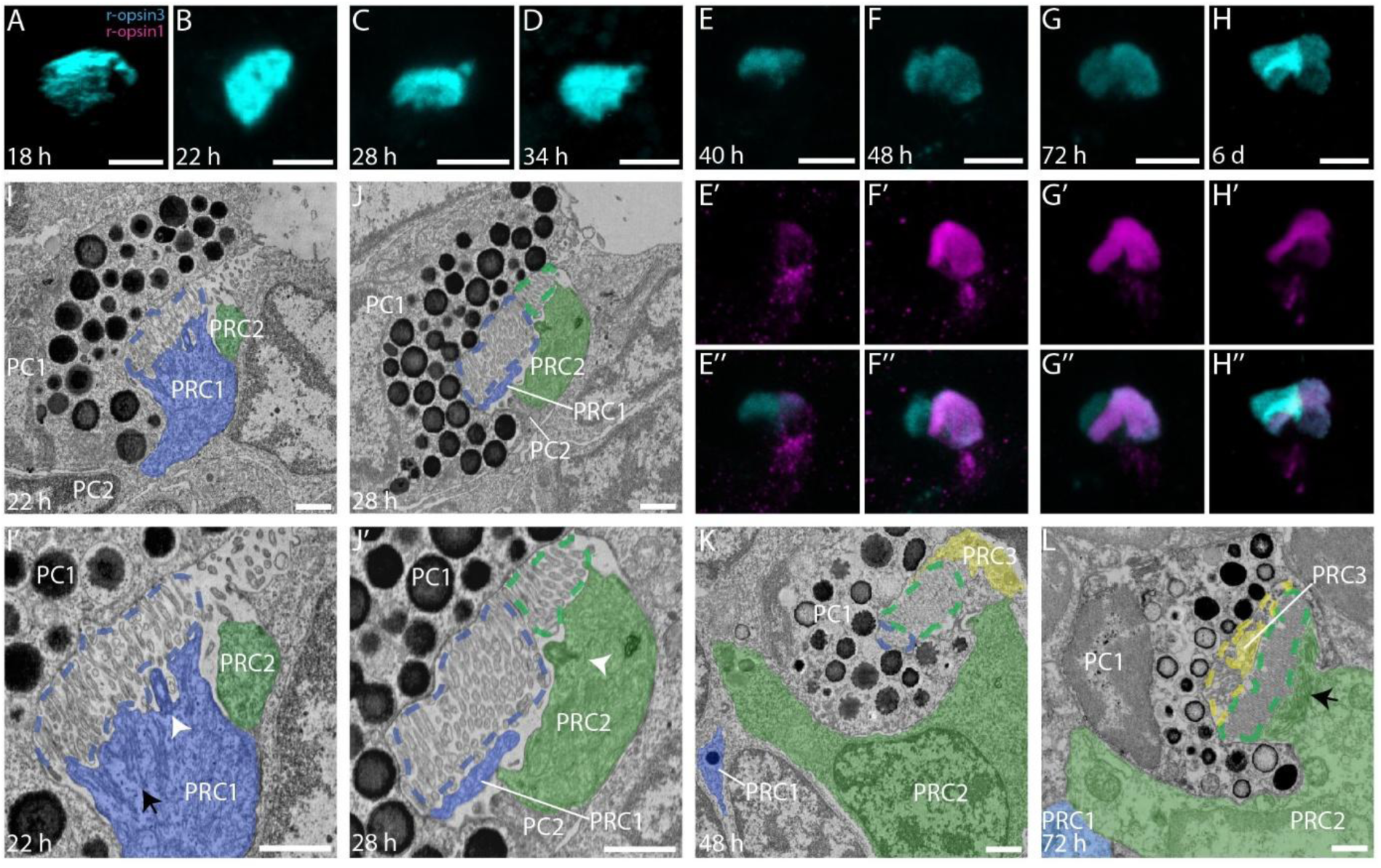
Organization of the right ventral eye. (A-H’’) Antibody staining of r-opsin3 (cyan) and r-opsin1 (magenta). (I-L) Overview of 22hpf PRC1, PRC2, PC1 and PC2, as well as detail showing PRC1 microvilli (blue dashed line), submicrovillar cisternae (black arrow) and cilium (white arrowhead). (J,J’) Overview and detail of 28hpf PRC1 and PRC2 microvilli (blue and green dashed line, respectively), as well as PRC2 cilium. (K) 48hpf PRC1, PRC2 and PRC3. (L) 72hpf PRC1-3, showing the PRC3 microvilli perpendicular orientation (yellow dashed line). PC = pigment cup, PRC = photoreceptor cell. Scale bars: A-H, 5 µm; I-L, 1 µm.

### (b) Organization of the eyespots

The larvae of *M. fuliginosus* develop three pairs of pigmented eyespots in the head, one mediodorsal, one laterodorsal, and one on a more ventral position which appear at different stages of development (Fig. 2; Sup.Fig.1). The ventral eyespots develop first at around 14 hpf, and the two dorsal eyespots form later, at around 42 hpf. While the ventral eye and the mediodorsal eyes exhibit a rhabdomeric structure, the dorsal lateral-most eyes have a ciliary structure. Both the ventral and mediodorsal eyespots have an inverted orientation, as microvilli located deep in the pigment cup receive light only after passing through the cell body (Fig. 3,5, Sup. Fig. 5-8).

**Figure 4.**
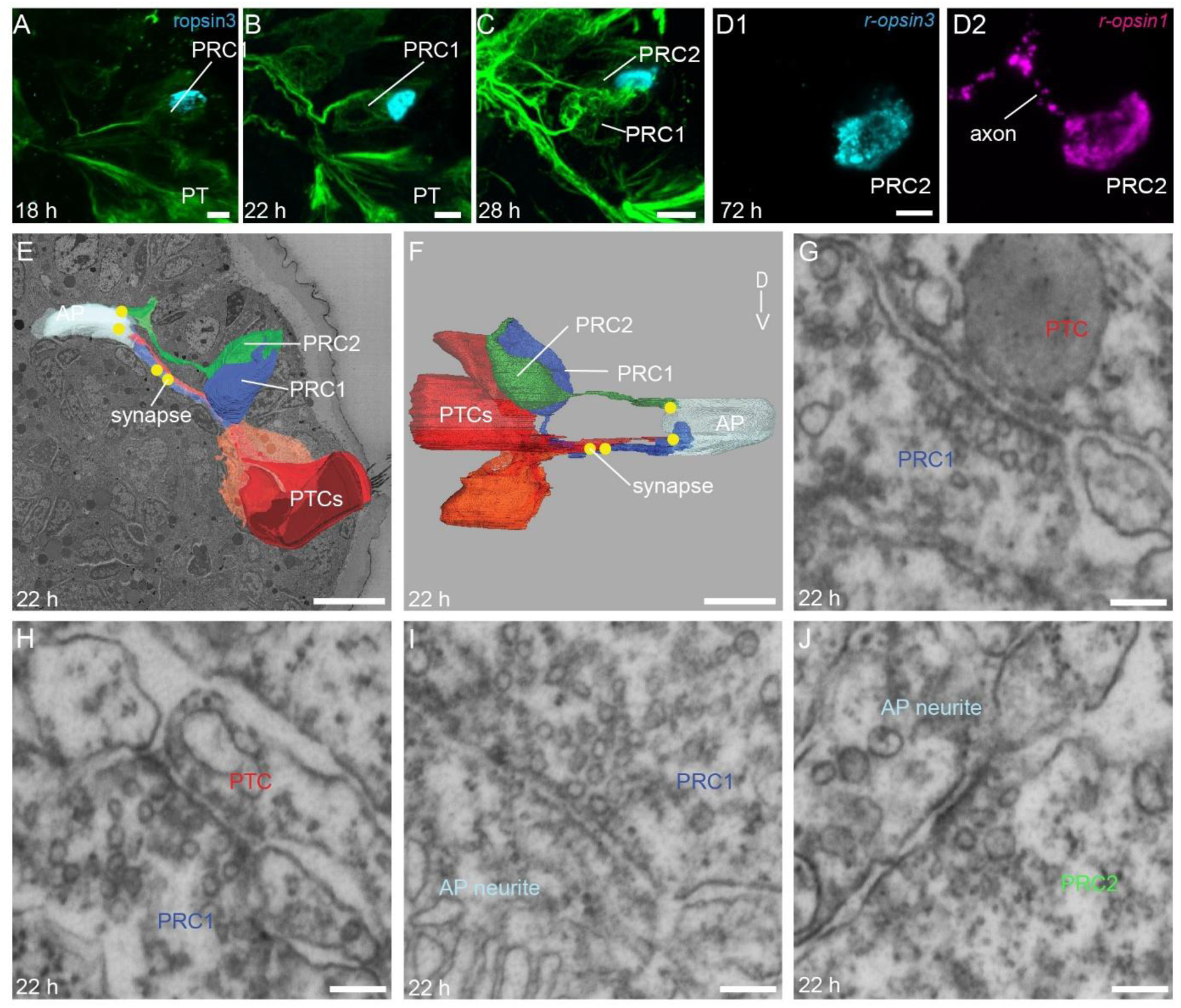
Right ventral eye and its circuitry. (A-C) Dorsal view of the eye and nervous system as revealed by r-opsin3 expression (blue) and acetylated α-tubulin immunostaining, green), respectively. (D1,D2) *In situ* hybridization at 72 h stage reveals that only r-opsin1 is present in the axon of PRC2. (E,F) Dorsal and anterior view on 3D reconstructions of PRC1, PRC2, prototroch cells, apical plexus and synapses (yellow dots). (G-J) Detail of synapses between PRC1 axon and PTC process (G,H) PRC1 axon and apical plexus neurite (I) and PRC2 axon and apical plexus neurite (J). AP = apical plexus, PRC = photoreceptor cell, PTC = prototroch cell. Scale bars: A-D, 5 µm; E,F, 10 µm; G-J, 100 nm.

**Figure 5:**
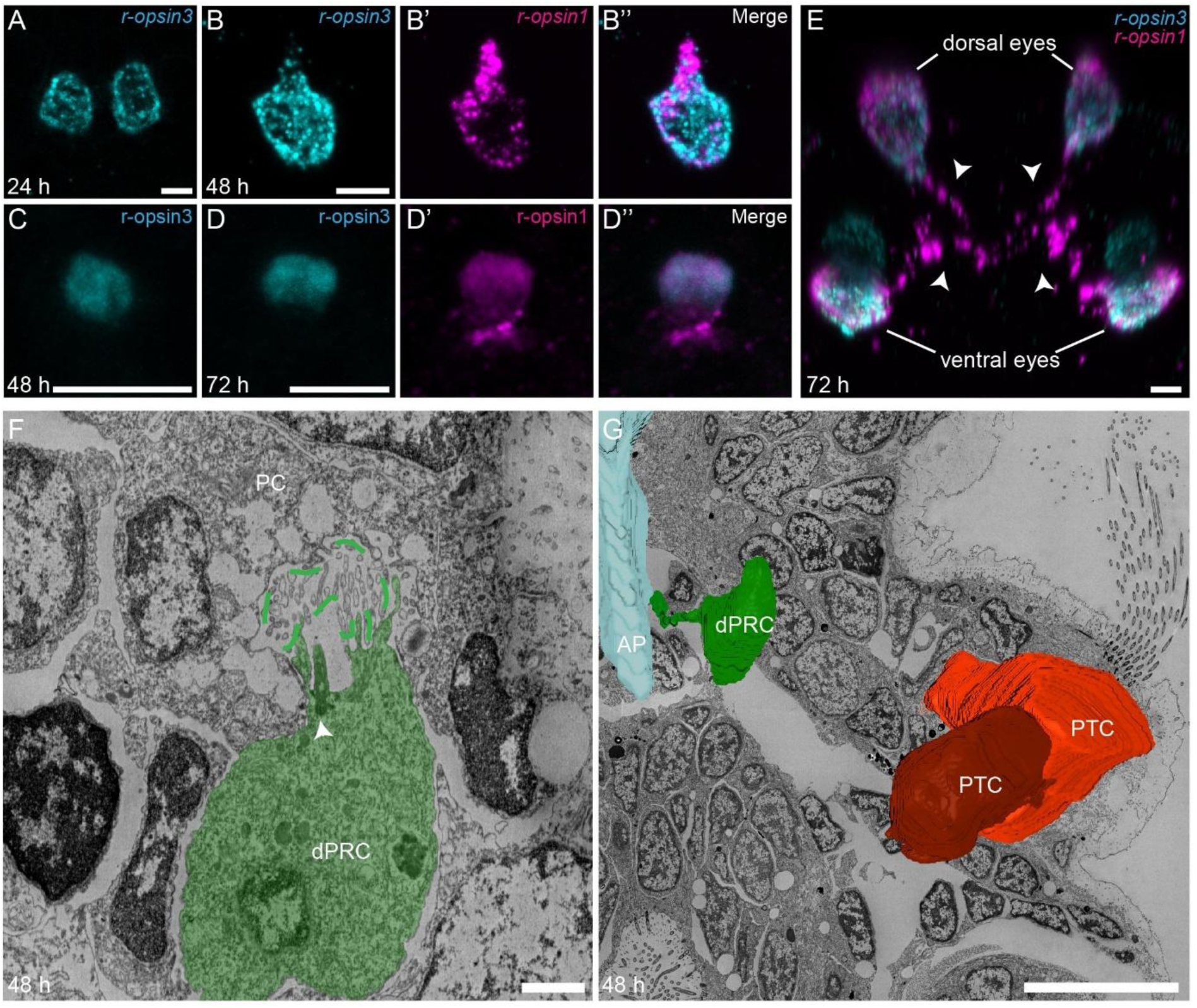
Dorsal eye and its circuitry. (A-B’’,E) Double FISH of r-opsin1 (magenta) and r-opsin3 (cyan) at 24 h (dorsal eye pair), 48 h (single dorsal eye) and 72 h (both ventral and dorsal eyes). The arrowheads in E indicate the axonal projections from the eyes. (C-D’’) Double immunostaining of r-opsins at 48 h and 72 h. (F) Dorsal eye PRC with cilium (arrowhead) and microvilli projecting into the pigment cup. (G) 3D reconstruction of the dorsal PRC, apical plexus and prototroch cells. AP = apical plexus, dPRC = dorsal photoreceptor cell, PC = pigment cup, PTC = prototroch cell. Scale bars: A-E, 5 µm; F, 1µm; G, 10 µm.

To enable a general comparison of annelid cerebral eyes, we therefore focused our analysis on the rhabdomeric eyes. We characterized their organization using 3D EM image stacks from early developmental stages: 22, 28, 48 and 72 hpf. The dorsal eyespots are structurally very simple, consisting of a single pigment cell and a single PRC extending numerous microvilli into the pigment cup (Fig. 5). The ventral eyes are slightly more complex and are positioned within the epidermis underneath the cuticle (Fig. 2). At 22 and 28hpf the ventral eye consists of two photoreceptor cells (PRC1 and 2), one pigment cell and one pigment cell precursor lying basal to the pigment cell (Fig. 2, 3; Sup. Fig. 5, 6). By 48 and 72 hpf, the ventral eye consists of three PRCs (PRC1-3), and two pigment cells. The PRCs are adjacent to each other and increase in size through the developmental stages. The PRC1 is located most posterior (22 and 28hpf) and dorsomedial (48 and 72 hpf) and PRC3 is most anterior and lateroventral (Fig. 2I-N’, 3).

The photoreceptor cells consist of an array of parallel microvilli, which are projected to the concavity of the pigment cell and parallel to the surface of the epithelium. Beneath the microvilli parallel submicrovillar cisternae and numerous mitochondria are present. At 22 and 28 hpf, the microvilli array of PRC2 lies lateral to that of PRC1 and is close to the epidermis (Fig.3I-J’; Sup. Fig. 5, 6). The microvillar array of PRC1 is larger than that of PRC2 at these early stages but decreases in size at 48 and 72 hpf. At this stage, PRC3 has the smallest microvilli array, which is projected to the innermost part of the pigment cup and perpendicular to the other microvilli (Fig. 3K, L; Sup. Fig. 7, 8). Basal bodies are present in all PRCs throughout all stages, as well as the pigment cells.

Based on the position and orientation of the three eyespot pigment cups, we estimate that these eyes provide wide viewing angles covering the dorsal, lateral, and ventral fields. The ventral rhabdomeric eyes are oriented towards the ventral and lateral direction, dorsal rhabdomeric eyes are oriented dorsally whereas the dorsal ciliary eyes are oriented posteriorly and laterally (data not shown). The ventral and mediodorsal eyes persist in juvenile stages and are clearly identifiable in adults, but the fate of the larval dorsolateral eyespots remains unclear.

### (c) Differential expression of r-opsin1 and r-opsin3 during larval stages

We analyzed the expression patterns of *Mfu-r-opsin1* and *Mfu-r-opsin3* in various larval stages by fluorescent *in situ* hybridization. We first detected *Mfu-r-opsin3* expression at approximately 14 hpf in a pair of cells in the region where ventral eyespots later form. These cells also express the neuronal marker *synaptotagmin1* (Kumar et al., 2020; Sup. Fig. 4). Adjacent to this cell, a reddish pigment cup becomes visible at around 19 hpf. By approximately 22 hpf, a second PRC expressing *Mfu-r-opsin3* appears adjacent to and slightly lateral to the first PRC (Fig. 2I). Shortly thereafter, at around 34 hpf, the second PRC begins to coexpress *Mfu-r-opsin1* (Fig. 2K). By 40 hpf, a third PRC expressing only *Mfu-r-opsin3* appears anterior and lateral to the other two PRCs (Fig. 2L). Gradually, the second PRC coexpressing the two opsins expands in size accompanied by the markedly stronger expression of *Mfu-r-opsin1* than that of *Mfu-r-opsin3* (Fig. 2L-N). This shift in expression is consistent with our RNA-seq analysis of stage-specific gene expression (data not shown). Notably, *Mfu-r-opsin1* mRNA was also detected within the axon extending from this second PRC (Fig. 2N; 4D’).

By correlating these observations with the EM data described above, it becomes clear that the first developing cell is the posterior dorsomedial PRC, the second is the largest cell in the middle and the third being the lateral and anterior cell sending microvilli into the base of the pigment cup. This mapping is also consistent with the immunodetection of opsin proteins using custom antibodies raised against Mfu r-opsin1 and Mfu-r-opsin3 (Fig. 3A-H’’). The second PRC starts expressing r-opsin1 protein at around 40 hpf (Fig. 3E-E’’) and both Mfu-r-opsin1 and Mfu-r-opsin3 proteins localize within the pigment cup in the region of the large rhabdom originating from the second PRC coexpressing *Mfu-r-opsin1* and *Mfu-r-opsin3* transcripts. Mfu-r-opsin1 protein is in addition localized outside the pigment cup (Fig. 3E’-H’) matching the position of the perikaryon of the second PRC. Mfu-r-opsin3 protein signal that does not overlap with Mfu-r-opsin1 protein matches the positions in the EM image stack of the smaller rhabdoms originating from the first and third PRCs expressing only r-opsin3 mRNA.

The dorsal eye can be traced from 24 hpf onwards based on the early onset of *Mfu-r-opsin3* expression in a pair of cells located medially on the dorsal side (Fig. 5A). A pigment cup is visible at a much later stage, around 42 hpf, by which time these medial eyespots have shifted from a slightly posterior position to the same transverse plane as the ventral eyespots. By 48 hpf, the dorsal eye PRC begins to coexpress the Mfu-r-opsin1 transcript but not the protein (Fig. 5B-B’’,C). The Mfu-r-opsin1 protein can be detected only around 72 hpf (Fig. 5D-D’’), and this coexpression pattern is maintained at later stages. As observed for the ventral eye PRC that coexpresses both opsins, we also detected Mfu-r-opsin1 within the axon of the dorsal PRC (Fig. 5E).

### (d) Ventral and dorsal eye circuitry

Preliminary information on the course of PRC axons could be retrieved from anti-acetylated tubulin labeling and opsin mRNA localization. From 17 hpf onwards, an axon extends from the first Mfu-r-opsin3 expressing PRC running towards the developing apical plexus underneath the apical tuft of the larva (Fig. 4A). This axon eventually reaches the larval brain at 22 hpf.

To better understand the ventral eye circuitry, we traced the axons of PRC1-3 through our 3D EM image stacks. At all stages PRC1 and PRC2 (but not PRC3) send axons towards the larval brain, with PRC2 reaching directly anteromedial towards the brain, and PRC1 first ventral and then anteromedial (Fig. 4E,F; Sup. Fig. 9, 10). At 22 and 28hpf the axons of PRC1, but not PRC2, are enriched/darkened by neuro vesicles and mitochondria. In both stages, the processes of the two most lateral ciliated cells, which form the prototroch (prototroch cell 3 and 4, Seybold et al. 2025), reach towards the apical plexus and are postsynaptic targets of PRC1, but not PRC2. Both PRC1 and PRC2 axons synapse to neurites in the apical plexus (Fig. 4G-J; Sup. Fig. 10-12). As we followed the axonal development in subsequent stages, the axons from both the ventral and the dorsal eye appear to converge towards the same region of the brain (Fig. 4, 5).

### (e) Eye PRCs differentially express two vesicular neurotransmitter transporters

Most PRCs use glutamate, acetylcholine, or histamine as their main neurotransmitter (Thoreson & Witkovsky 1999; Farca-Luna & Sprecher 2013; Randel et al., 2014). Neurotransmitter identity is functionally defined by vesicular neurotransmitter transporters, which package specific neurotransmitters into synaptic vesicles and thereby mediate synaptic signaling.

To identify the neurotransmitter types used by PRCs in *M. fuliginosus*, we examined the expression of vesicular glutamate transporter (VGluT) and vesicular acetylcholine transporter (VAChT) as molecular markers. At 24 hpf, the single r-opsin3+ PRC in the ventral eye coexpresses both *Mfu-VAChT* and *Mfu-VGluT* (Fig. 6A, D). In contrast, the second and third PRCs of the ventral eye lack *Mfu-VAChT* (Fig. 6B,C1,C2) and express only the glutamatergic marker, *Mfu-VGluT* (Fig. 6E1,E2). Although both transporters are present in various domains of the brain, their coexpression is restricted to the first r-opsin3+ PRC.

**Figure 6:**
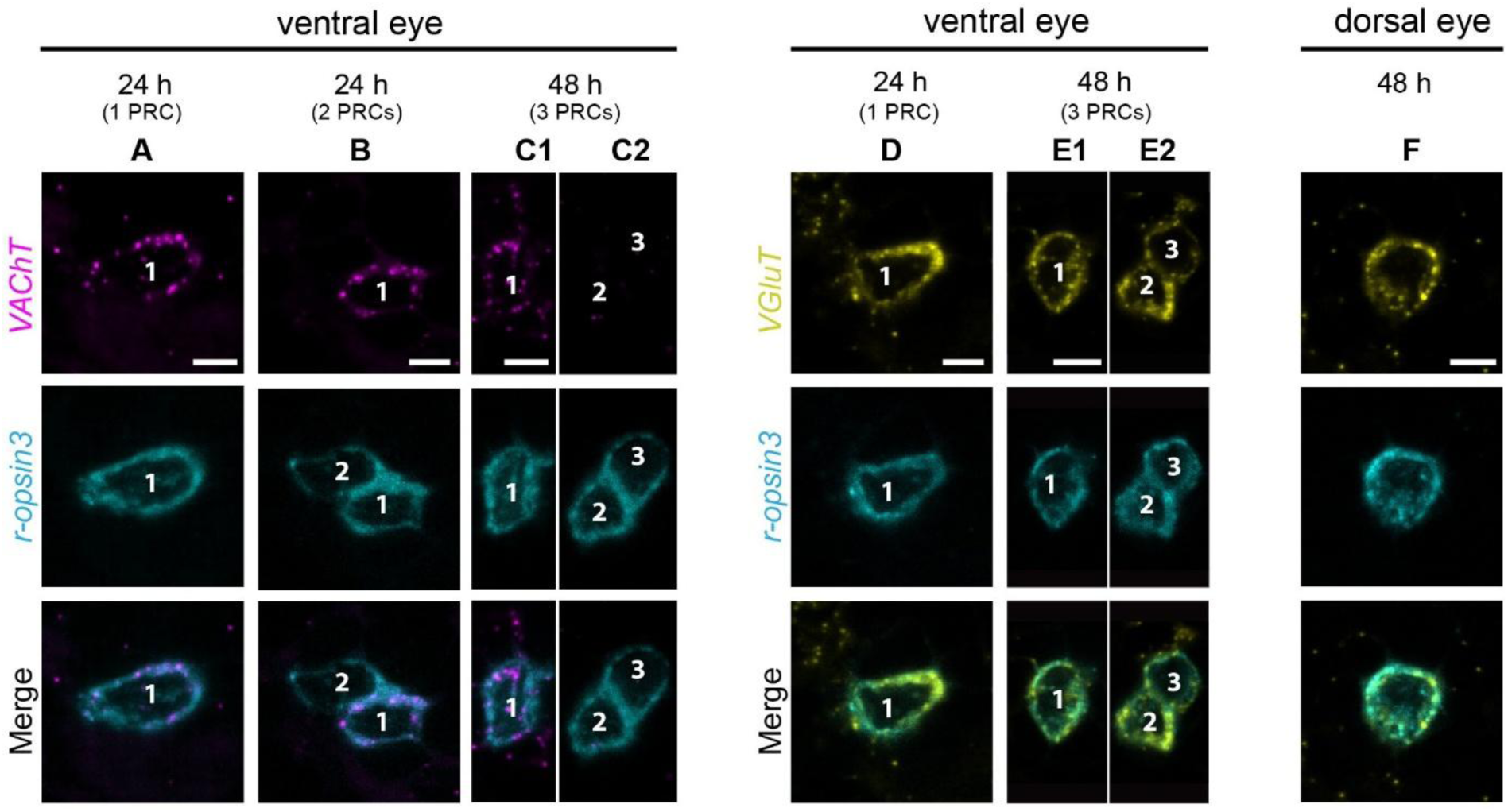
Expression of eye neurotransmitter receptors. Double FISH of neurotransmitter receptors, VAChT (magenta) or VGluT (yellow) with ropsin3 (cyan). Panels A-E2 are the ventral eye PRCs and the numbers indicate the developing order of the PRCs. At the 24 h stage, the second developing PRC is occasionally detectable (shown in B). Panel F is the dorsal eye PRC. The images are z projection of the eye region. Scale bar: 5 µm.

The dorsal eye PRC also expresses only *Mfu-VGluT* (Fig. 6F). Notably, the expression of *Mfu-VGluT* is much stronger than *Mfu-VAChT* within the PRCs. In later stages of development, the expression of *Mfu-VAChT* in the PRC gets much weaker (Fig. 6A-C1), while its expression expands in other brain regions (data not shown). In relation, *Mfu-VGluT* remains strongly expressed in both ventral and dorsal eye PRCs (Fig. 6D-F) and is additionally localized to smaller expression domains in the brain and apical region (data not shown).

### (f) Conserved set of genes specify the rhabdomeric eyes

Eye development across metazoans is known to depend on a conserved set of transcription factors, the core of which constitutes the Pax-Six-Eya-Dach regulatory network (Kumar 2001; Kozmik et al., 2007). To explore the role of these genes in the rhabdomeric eyes of *M. fuliginosus*, we analyzed the expression profiles of these candidate genes, together with *Mfu-Prox1* and *Mfu-Otx,* both important for eye specification. Expression analyses were initiated at the onset of *Mfu-ropsin3* expression in the ventral and dorsal eyes.

At 14 hpf, *Mfu-Pax6*, *Mfu-Otx*, and *Mfu-Six1/2* are strongly expressed in the region of the ventral eye. Colocalization with *Mfu-r-opsin3* confirmed their expression in the first ventral eye PRC (Fig. 7). At this stage, *Mfu-Prox1* shows weak expression in the PRC, but is strongly expressed in adjacent cells, which likely correspond to subsequently differentiating PRCs and pigment cells (Fig. 7). In contrast, *Mfu-Eya* is only weakly expressed in the PRCs, although its other expression domains largely overlap with its transcriptional co-activator, *Mfu-Six1/2* (Fig. 7).

**Figure 7:**
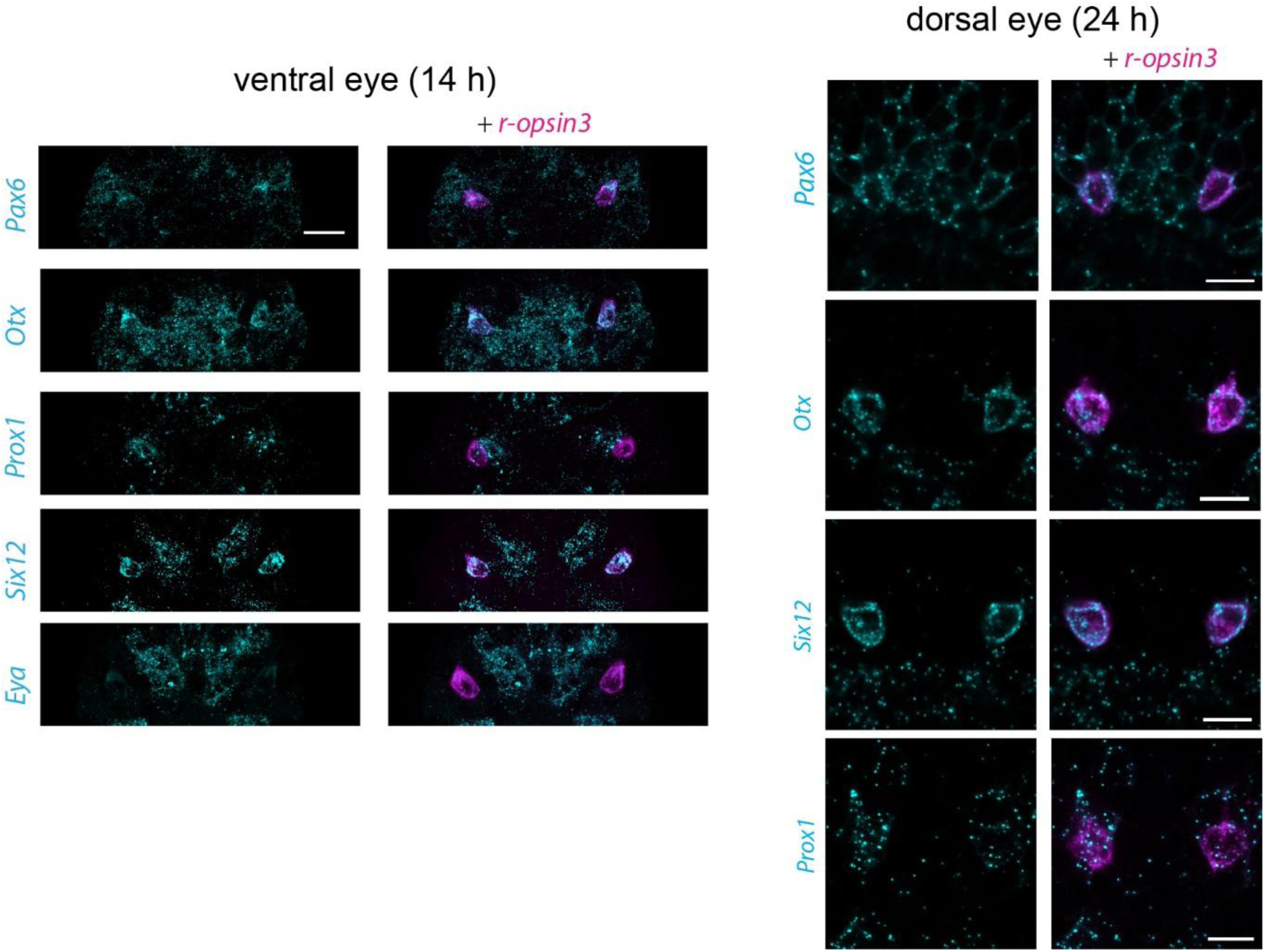
Expression of eye development genes in the ventral eye at 14 hpf and dorsal eyes at 24 hpf. Double FISH of eye genes and r-opsin3. The images are confocal z stacks of the ventral and dorsal eye region. Scale bars: 10 µm.

A similar pattern of expression is maintained at 24 hpf (data not shown), with all examined genes also expressed in the developing dorsal eye PRC (Fig. 7). By 48 hpf, these genes continue to be distinctly expressed in the PRCs of both ventral and dorsal eye. At this stage, in the ventral eye, *Mfu-Otx* is more strongly expressed in the larger cell coexpressing *Mfu-r-opsin1* and *Mfu-r-opsin3* while its expression is weaker in the other two PRCs. In contrast, the expression of *Mfu-Prox1* gets considerably weaker within the PRCs in both ventral and dorsal eyes (data not shown).

## Discussion

Annelids exhibit remarkable diversity in eye structures, suggesting multiple convergent and independent evolutions. However, the extent of conservation and innovation remains unclear due to limited comparative studies. Our study provides new insights into the molecular, ultrastructural, and circuit-level organization of cerebral eyes in the sedentary annelid *Malacoceros fuliginosus*. Comparison with the errant annelid *Platynereis dumerilii* reveals strong evidence of homology between their rhabdomeric cerebral eyes, revealing the ancestral annelid condition. Eye connectivity data from both ventral larval and dorsal adult eyes, together with molecular profiles, support this homology, reinforced by similar spatial and temporal *Mfu-r-opsin1* and *Mfu-r-opsin3* expression.

### (a) Differential expression of r-opsin1 and r-opsin3

The two canonical r-opsin paralogs in larval *M. fuliginosus* are restricted to the paired ventral and dorsal microvillar eyes and are differentially expressed in PRCs during development. *Mfu-r-opsin3* is the first opsin expressed in all PRCs, whereas *Mfu-r-opsin1* appears later, only in PRC2 of the ventral eye and in the dorsal PRC. Therefore, both eye pairs exhibit a temporal offset in opsin deployment, with r-opsin3 expression preceding r-opsin1 at both transcript and protein levels. In *Capitella teleta*, only a single r-opsin orthologous to *Mfu-r-opsin3* is expressed in the larval eye, and its mutation prevents phototaxis (Neal et al., 2019).

PRC2 enlarges during development, with an increasing *Mfu-r-opsin1/Mfu-r-opsin3* expression ratio. This suggests *Mfu-r-opsin3* functions in early larval photoreception, while *Mfu-r-opsin1* dominates later. Spectral sensitivity or molecular differences between the paralogs remain unknown, though opsin coexpression may enhance sensitivity rather than discrimination (Dalton et al., 2014; Gühmann et al., 2015).

### (b) Comparison of *M. fuliginosus* and *P. dumerilii* microvillar eyes

Our phylogeny provides clear evidence that *Mfu-r-opsin1* and *Mfu-r-opsin3* are orthologous to *Pdu-r-opsin1* and *Pdu-r-opsin3*. These paralogs diverged in the last common ancestor of Errantia (*P. dumerilii*) and Sedentaria (*M. fuliginosus*), making their comparison informative for eye evolution.

The ventral larval eye of *P. dumerilii* has a simple few-celled inverse organization without accessory structures (Rhode, 1992; Randel et al., 2013), similar to *M. fuliginosus*. *P. dumerilii* has two PRCs and a pigment cup until 72 hpf, whereas *M. fuliginosus* has three PRCs and two pigment cup cells in late larval stages. All PRCs have prominent microvilli with rudimentary cilia; the PRC2 cilium in *M. fuliginosus* is more distinct, and all PRCs express *Xenopsin* (Vöcking et al., 2017; Döring et al., 2020) but this opsin type is secondarily lost in *P. dumerilii*. Pigment cups in both species orient ventrally and posteriorly, shaping similar light incidence angles.

Developmental opsin expression follows a conserved sequence: the first PRC expresses only r-opsin3, and the second coexpresses r-opsin3 and r-opsin1 (Jékely et al., 2008; Randel et al., 2013). In *P. dumerilii*, r-opsin3 declines in the ventral larval eyes, while in *M. fuliginosus*, it persists, with r-opsin1 gradually dominating (ratio 1:1 at 48 hpf to 10:1 at 10 dpf). Notably, in both species the temporal offset of r-opsin1 deployment is conserved. A third PRC expressing r-opsin3 develops in *M. fuliginosus*; a similar PRC in *P. dumerilii* is not reported. These data indicate a conserved order in PRC development and opsin deployment.

At the ultrastructural level the dorsal eyes differ markedly. In *M. fuliginosus*, the dorsal eye is a simple inverse structure with a single PRC and pigment cell until 10 dpf, remaining small throughout life. *P. dumerilii* develops large dorsal eyes with hundreds of PRCs and pigment cells, adopting an everse organization. Early development is comparable, but later stages diverge. Although both species coexpress r-opsin1 and r-opsin3, the precise temporal dynamics of r-opsin1 deployment have not been explicitly described in *P. dumerilii*. This likely reflects the broad developmental sampling in previous studies which may have obscured the temporal offset of r-opsin1 in the dorsal eye, similar to those observed for ventral eye PRC2. The dorsal eye PRC in *M. fuliginosus* also has a rudimentary cilium and expresses *Xenopsin* (Vöcking et al., 2017; Döring et al., 2020)

### (c) Neuronal projection of the eyes and neurotransmitter expression

In *P. dumerilii*, the first ventral eye PRC functions during early development, before maturation of the brain and neuronal scaffold. Its axon initially projects laterally to the prototroch, joins the prototroch ring nerve, and forms cholinergic synapses with prototroch cells, enabling brain-independent phototactic steering (Jékely et al., 2008), before later turning toward the developing brain (Jékely et al., 2008). At later stages, phototactic steering is mediated by the adult dorsal eye, whose glutamatergic PRCs project to brain interneurons that regulate ciliary activity and musculature via a more complex circuit (Randel et al. 2014, 2015), similar to *M. fuliginosus*.

In *M. fuliginosus*, the ventral eye can likewise mediate phototaxis at the single-PRC stage. PRC1 extends an axon early (∼16 hpf), prior to brain and scaffold maturation (Seybold et al., 2025), projecting medially past prototroch basal extensions before turning anteriorly toward the prospective brain. Along this trajectory, PRC1 forms synapses with prototroch cell extensions, acting as a sensory-motor neuron analogous to the first ventral eye PRC of *P. dumerilii*. Only PRC1 synapses with prototroch cells, whereas PRC1 and PRC2 synapse with neurites in the apical plexus, as described in *P. dumerilii*, where they connect to motor neurons via distinct circuits (Randel et al. 2013, 2015). Ventral PRC3 and dorsal PRC axons also extend toward the brain but are shorter and lack synapses, indicating ongoing differentiation.

The first PRC in *P. dumerilii* is reported to be exclusively cholinergic (Jékely et al. 2008; Randel et al. 2014), while the transmitter identity of the second and third PRCs remains unknown. In contrast, the corresponding PRC in *M. fuliginosus* coexpresses VAChT and VGluT, whereas all other ventral eye PRCs express VGluT exclusively. The expression of dual transmitter in the first PRC suggests temporal and spatial dimensionality with acetylcholine contributing to sensory-motor signaling and the subsequent glutamatergic connection mediating brain-projecting visual circuit.

### (d) Conserved gene networks in annelid eyes

As in bilaterian cerebral eyes, annelid eye development is influenced by a conserved gene set (Arendt et al., 2002; Vopalensky & Kozmik 2009; Gehring 2014). Pax6, Otx, Six1/2 and Prox1 are involved in specification of both ventral and dorsal microvillar eyes, consistent with findings from other protostome cerebral microvillar eyes (Martín-Durán et al. 2012; Samadi et al. 2015; Vöcking et al. 2015; Zhu et al. 2017). Lineage-specific divergence is indicated by weak or absent expression of *eya* and *dach*, which are essential in other protostomes (Shen & Mardon 1997; Yang et al. 2009; Jin & Mardon 2016), suggesting either very low or early expression, or a reduced role in eye specification in *M. fuliginosus*. Nevertheless, the presence of a conserved transcription factor network likely regulating both rhabdomeric eyes supports their homology to cerebral eyes of other Bilateria.

### (e) Evolutionary origin of ventral and dorsal eyes

Shared position, developmental onset, structural features, and gene expression indicate a common origin of ventral eyes in *P. dumerilii* and *M. fuliginosus*. The expression of r-opsin3 and cholinergic marker in the first PRC and the conserved role in larval phototaxis further supports homology. The dorsal eyes, despite structural differences (everse in *P. dumerilii*, simple inverse in *M. fuliginosus*), likely share a common evolutionary origin. Comparative studies in errant polychaetes suggest adult eye complexity evolved stepwise (Suschenko & Purschke, 2009; Purschke & Nowak, 2015). Few sedentary polychaetes have multicellular everse eyes (e.g., *Scoloplos armiger*, *Flabelligera affinis*), while most show simple inverse organization (Purschke et al., 2006; Wilkens & Purschke 2009; Vodopyanov & Purschke 2017). Shared position, later onset, r-opsin coexpression, VGluT expression, and brain projections support dorsal eye homology.

Homology implies at least two eye pairs in the last common ancestor of Errantia and Sedentaria. Data from basally branching annelids are limited: *Owenia fusiformis* has late larval eyes retained post-metamorphosis, absent in *Myriowenia* (Helm, 2016; Beckers, 2019; Carrillo, 2024); *Magelona* sp. have transient larval eyes (Wilson, 1982); chaetopterids have two pairs of larval/adult eyes (Irvine et al., 1999; Helm et al., 2022; Purschke et al., 2022); *Amphinomida* possess two pairs of adult rhabdomeric eyes (Purschke et al., 2022). Sipunculan larvae show multiple eyespots (Rice, 1978; Radashevsky, 2006). This diversity complicates comparisons and highlights the need for additional molecular and developmental studies.

R-opsin duplication likely preceded eye duplication, as suggested by the conserved differential r-opsin expression in *M. fuliginosus* and *P. dumerilii*. The evolution of PRC signaling, whether acetylcholinergic sensory–motor or glutamatergic brain-projecting, remains open, although early acetylcholine signaling may represent a common annelid trait.

### Conclusion and future perspectives

Closely related species are ideal for studying sensory evolution and adaptation. Despite annelid eye diversity, the last common ancestor of Errantia and Sedentaria likely had two pairs of simple microvillar eyes. Complex multicellular eyes arose via the addition of sensory and pigment cells and accessory structures. Opsin gene duplication, especially r-opsin1 and r-opsin3, contributed to functional diversification, with r-opsin3 mediating early phototaxis and r-opsin1 acting at later stages (Jékely et al., 2008; Neal et al., 2019; Giacinto Vivo et al., 2023). Further studies on molecular physiology and regulation of r-opsins will clarify adaptive significance.

Broader ultrastructural, gene regulatory, and opsin expression analyses in basal annelids, coupled with eye circuitry studies at synaptic resolution, will deepen understanding of eye evolution. Comparative larval studies in lophotrochozoans can reveal whether sensory-motor circuitry is ancestral or annelid-specific, aiding phototactic behavior before CNS maturation.

## Supplementary figures

**Sup. Fig. 1:**
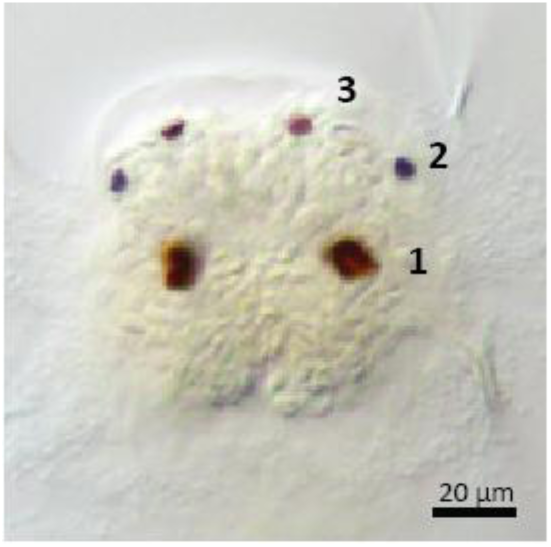
Front view of 48hpf *M. fuliginosus* larvae showing the eyes with the numbers indicating the order of their appearance.

**Sup. Fig. 2:**
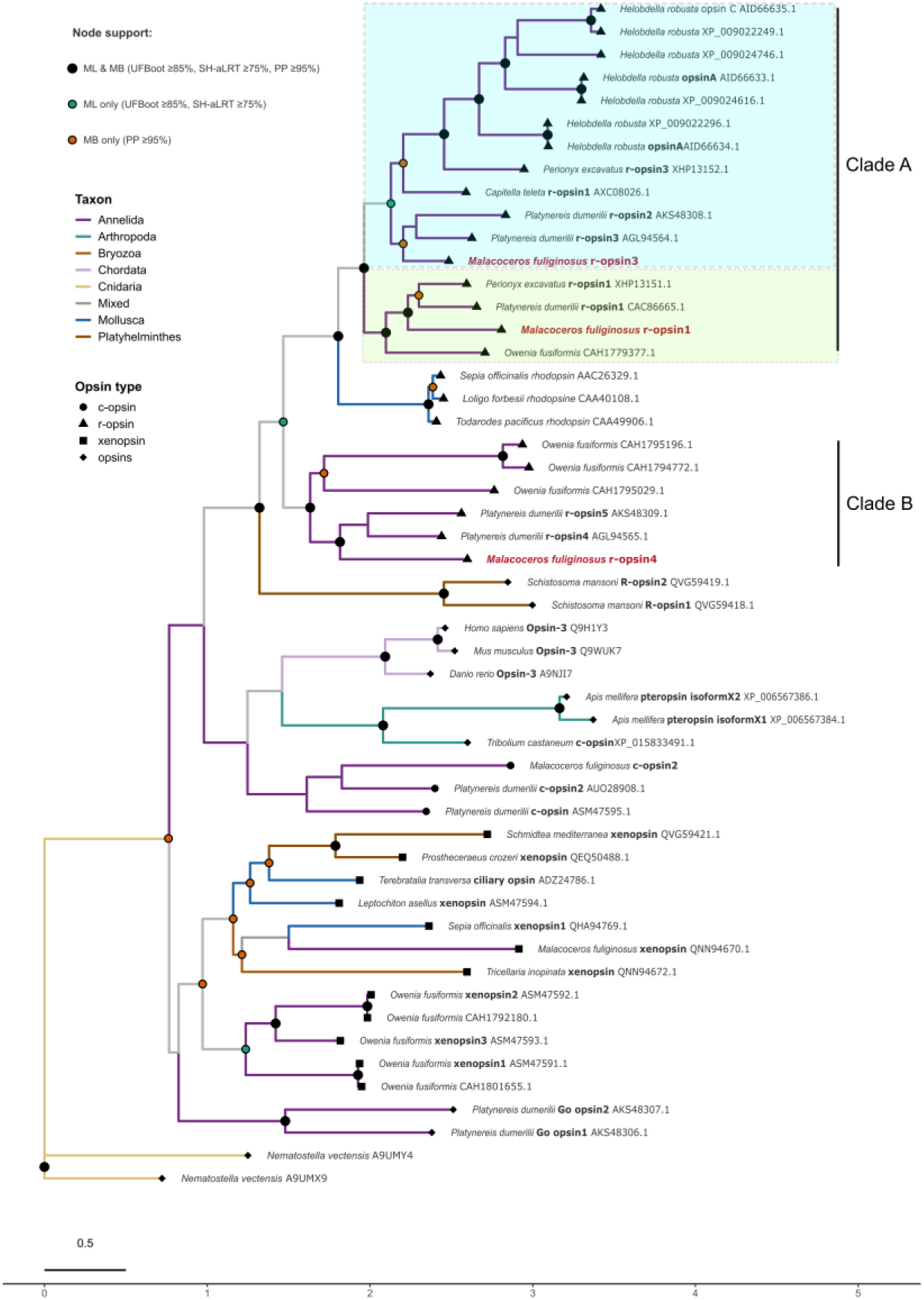
Phylogenetic tree of opsins.

**Sup. Fig. 3:**
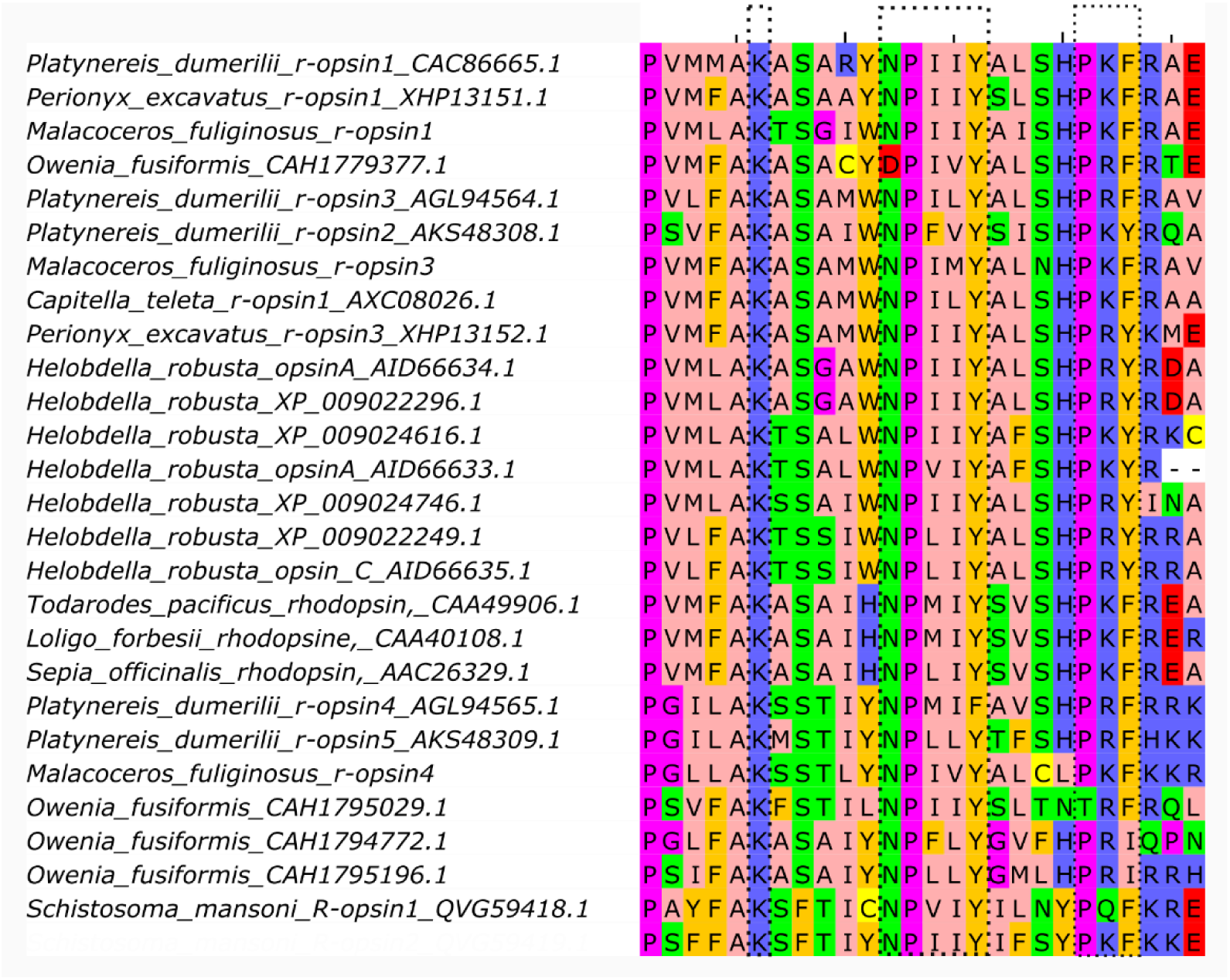
Conserved squence motifs of r-opsins

**Sup. Fig. 4:**
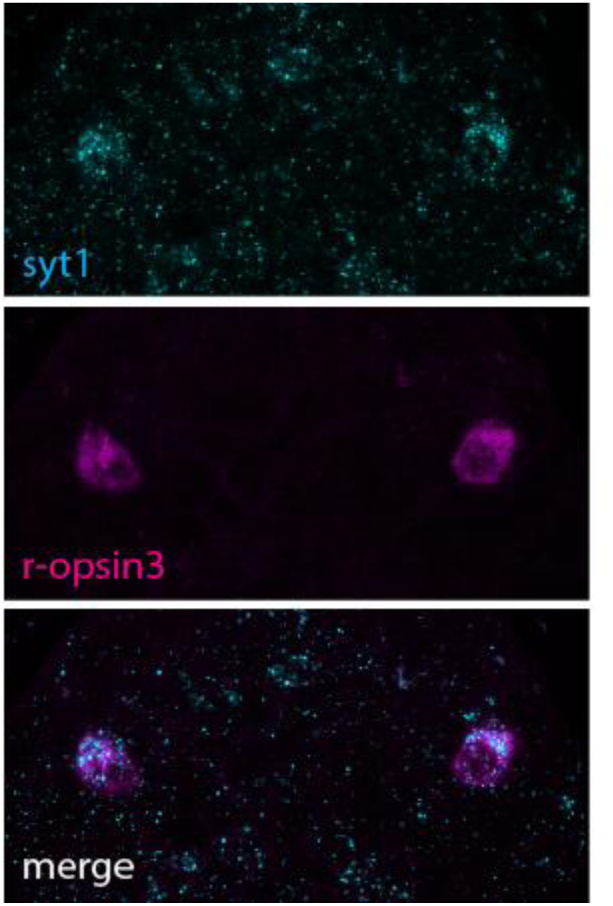
Double FISH of the *Mfu-synaptotagmin1* and *Mfu-r-opsin3* at 14 hpf.

**Sup. Fig. 5:**
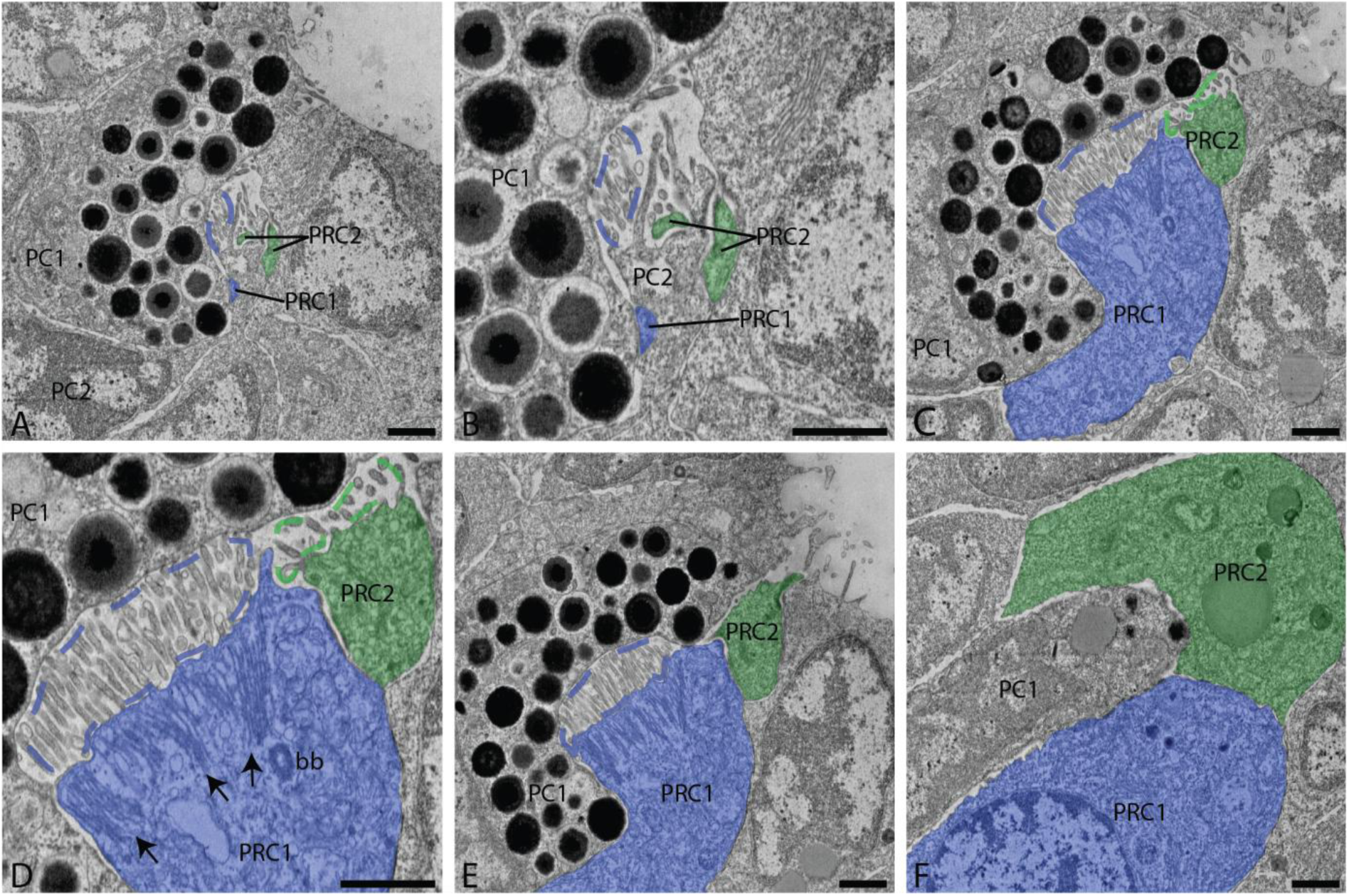
Ultrastructural organization of the right ventral eye at 22hpf from dorsal to ventral. (A,B) Overview and detail of PCR1, PRC2 PC1 and PC2. (C,D) Overview and detail of PRC1 and PRC2 microvilli (blue and green dashed line, respectively), as well as PRC1 submicrovillar cisternae (black arrows) and basal body (bb). (E) Apical extension of PRC2. (F) Nuclei of PRC1 and PRC2. PC = pigment cup, PRC = photoreceptor cell. Scale bar: 1 µm.

**Sup. Fig. 6:**
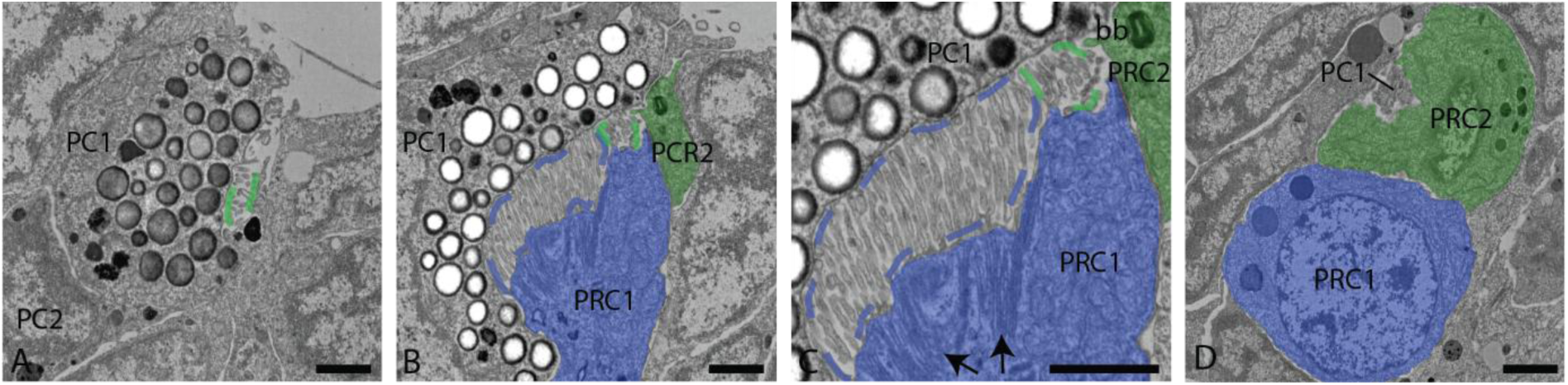
Ultrastructural organization of the right ventral eye at 28hpf from dorsal to ventral. (A) PRC2 microvilli (green dashed line) projecting into PC1. (B,C) Overview and detail of PRC1 (blue dashed line) and PRC2 microvilli, as well as PRC1 submicrovillar cisternae (black arrows) and PRC2 basal body (bb). (D) Nuclei of PRC1 and PRC2. PC = pigment cup, PRC = photoreceptor cell. Scale bar: 1 µm.

**Sup. Fig. 7:**
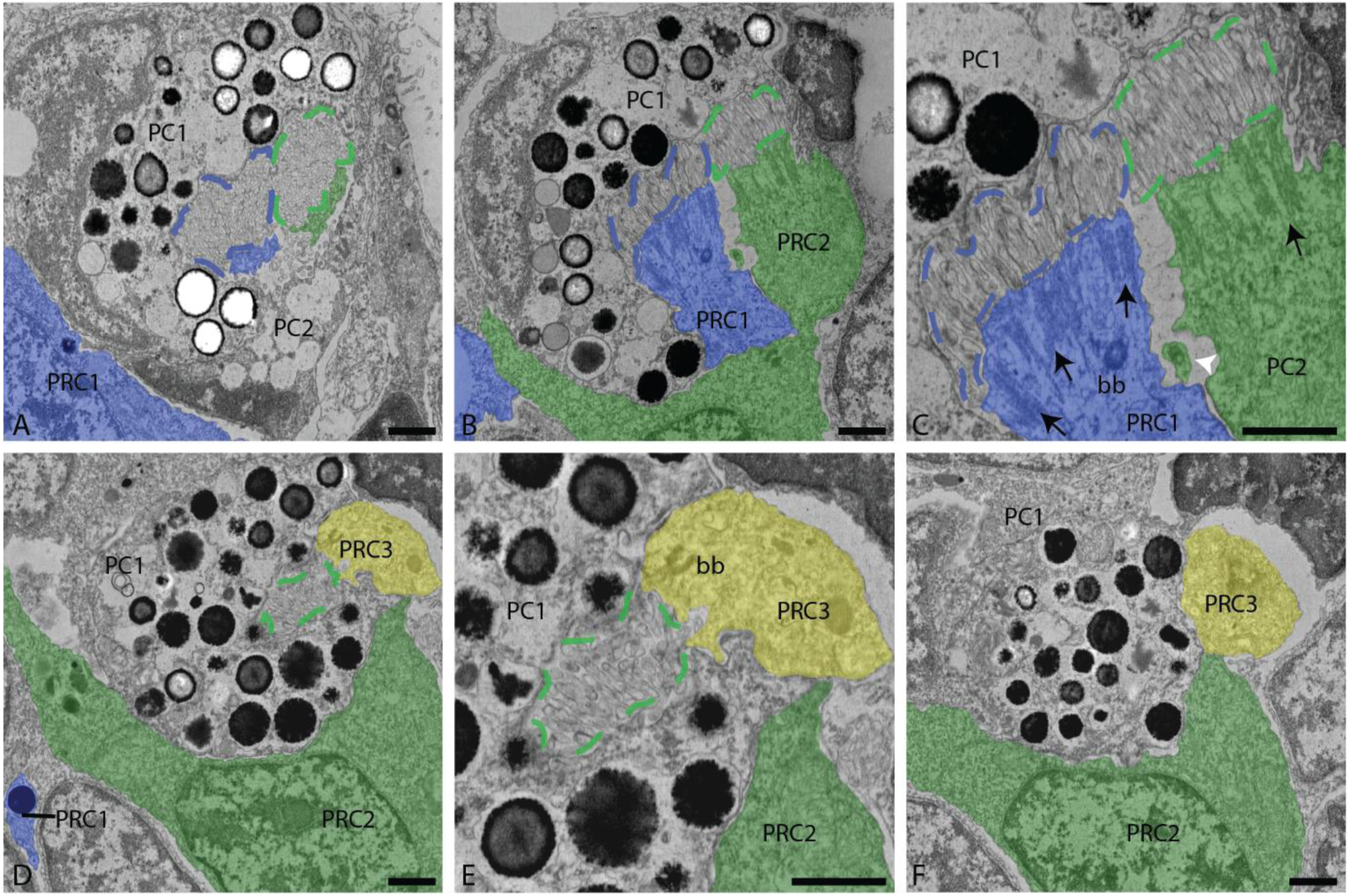
Ultrastructure of the right ventral eye at 48hpf from dorsal to ventral. (A) PC1 and PC2 surrounding PRC1 and PRC2 microvilli (blue and green dashed line, respectively). (B,C) Overview and detail of PRC1 and PRC2 microvilli and submicrovillar cisternae (black arrows), as well as PRC1 basal body (bb) and PRC2 cilium (white arrow head). (D,E) Overview and detail of PRC3 with basal body. (F) PRC3 nucleus. PC = pigment cup, PRC = photoreceptor cell. Scale bar: 1 µm.

**Sup. Fig. 8:**
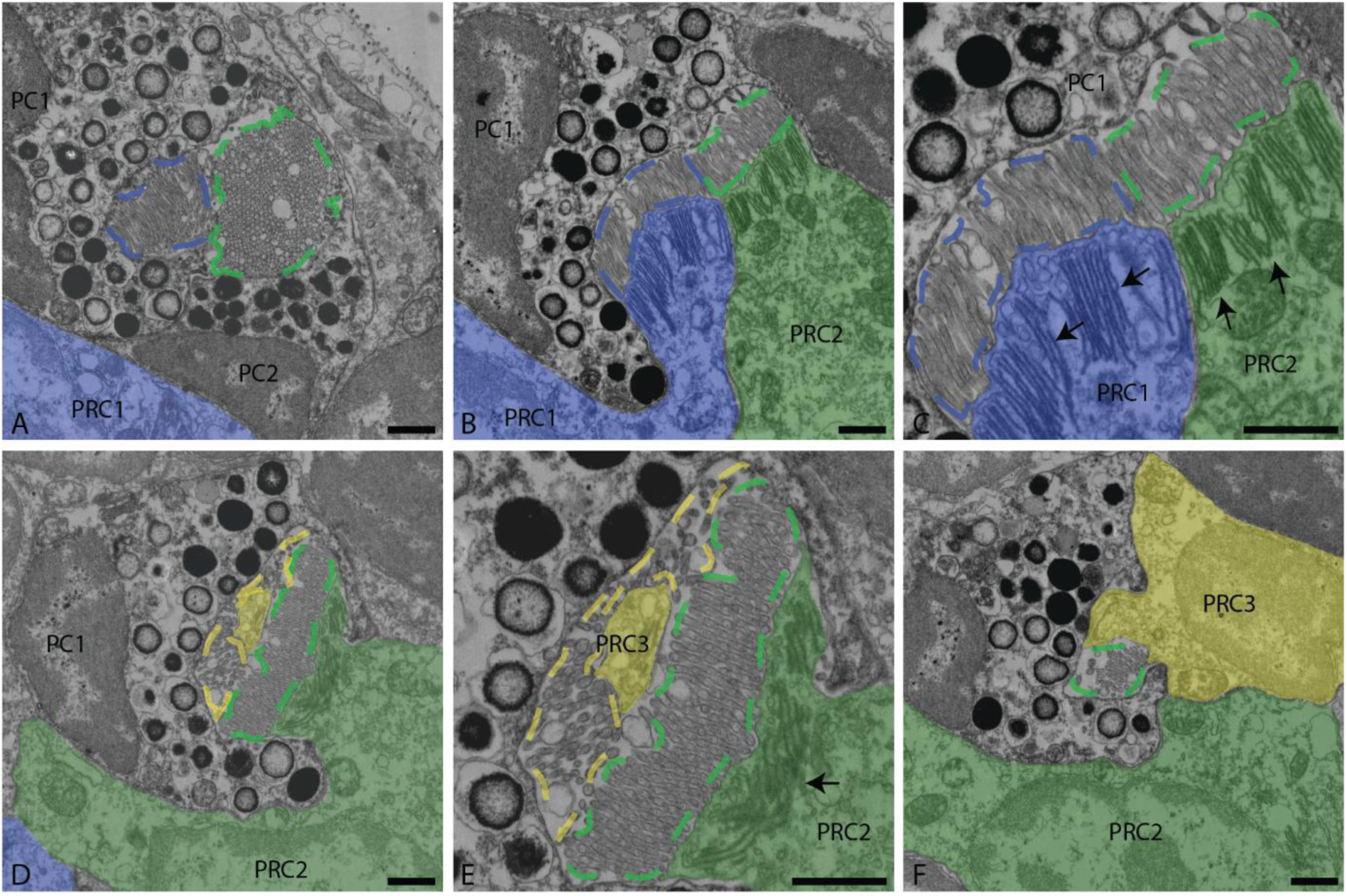
Ultrastructure of the right ventral eye at 72hpf from dorsal to ventral. A) PC1 and PC2 surrounding PRC1 and PRC2 microvilli (blue and green dashed line, respectively). B,C) Overview and detail of PRC1 and PRC2 microvilli, as well as submicrovillar cisternae (black arrows). (D+E) Overview and detail of PRC3 in the innermost part of the pigment cup and perpendicular to the PRC2 microvilli. (F) PRC3 nucleus. PC = pigment cup, PRC = photoreceptor cell. Scale bar: 1 µm.

**Sup. Fig. 9:**
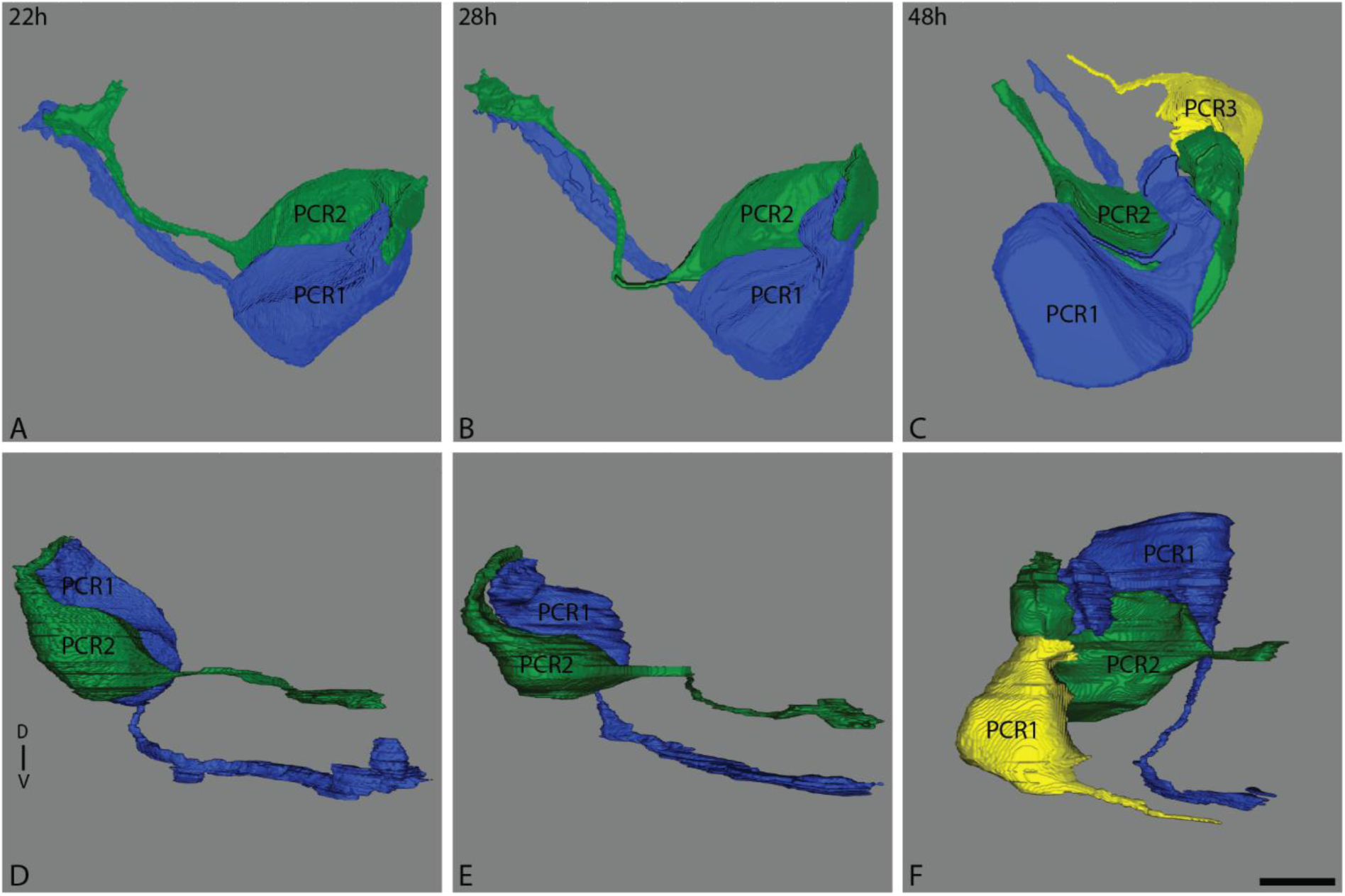
3D reconstruction of PRC1, PRC2 and PRC3 viewed from dorsal (A-C) and from anterior (D-F) at 22, 28 and 48hpf. Scale bar, 5µm.

**Sup. Fig. 10:**
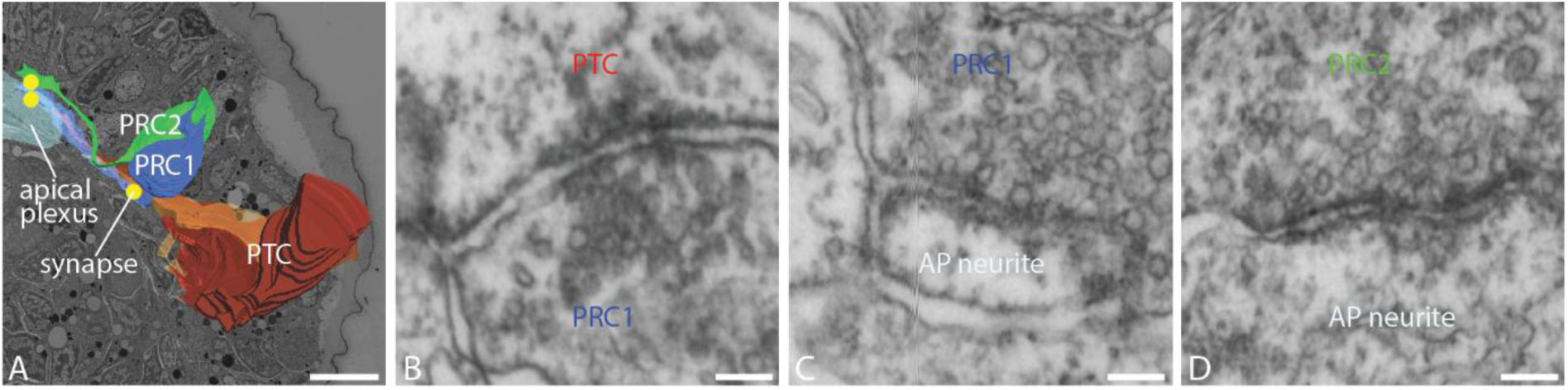
Ultrastructural organization of the right ventral eye and its circuitry at 28hpf. A) dorsal view on 3D reconstructions of PRCs, prototroch cells, apical plexus and selected synapses (yellow dots). B-D) Detail of selected synapses between PRC1 axon and PTC process (B), PRC1 axon and AP neurite (C) and PRC2 axon and AP neurite (D). AP = apical plexus, PRC = photoreceptor cell, PTC = prototroch cell. Scale bars: A, 10 µm; B-D, 100 nm.

**Sup. Fig. 11:**
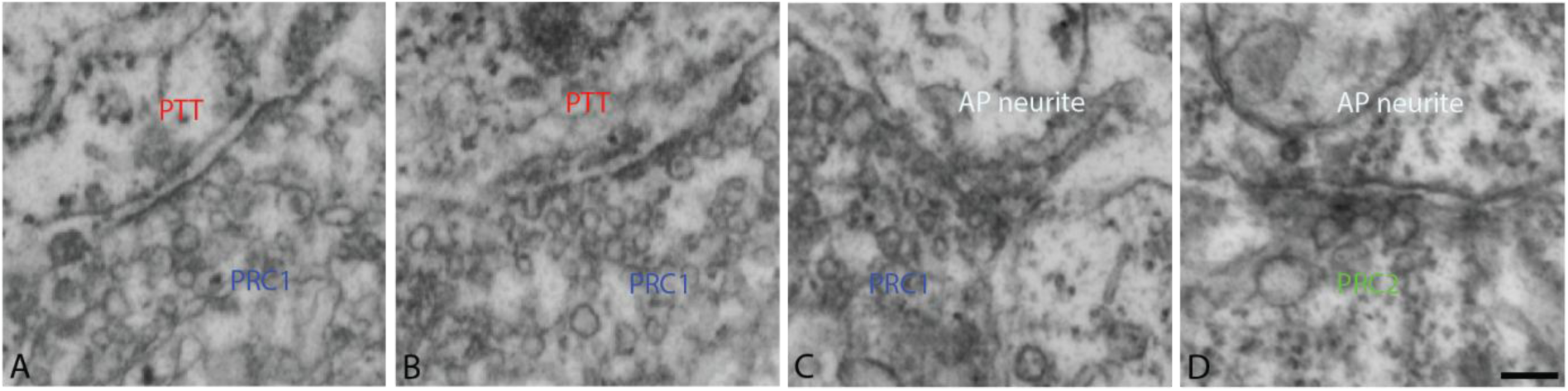
Synapses of the ventral eye at 22hpf. Synapses between PRC1 axon and PTT process (A,B), PRC1 axon and AP neurite (C) and PRC2 axon and AP neurite (D). AP = apical plexus, PRC = photoreceptor cell, PTC = prototroch cell). Scale bar: 100 nm.

**Sup. Fig. 12:**
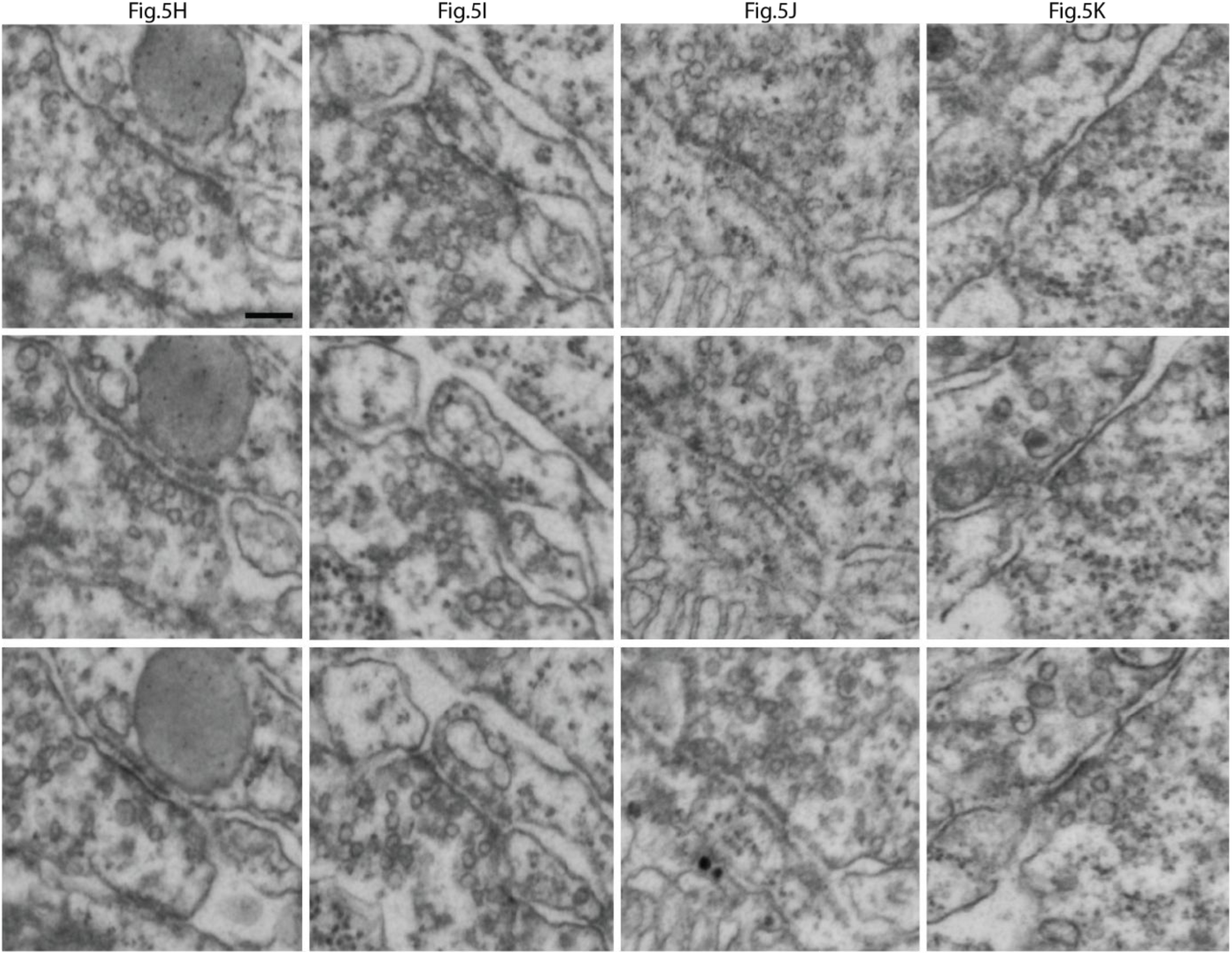
Series of sections through synapses displayed in Fig.5. H-K. Scale bar, 100 nm

